# Resurrected in the field: benefits of rapid adaptation to historic drought seen mainly at the leading edge of a plant species’ range

**DOI:** 10.64898/2026.03.12.711156

**Authors:** Lillie K. Pennington, Jason P. Sexton

**Affiliations:** Department of Life and Environmental Sciences, University of California, Merced, Merced, USA; Genetics Department, University of Georgia, Athens, USA

**Keywords:** rapid adaptation, resurrection study, leading edge, rear edge, range limits, native plant, field study, common garden

## Abstract

Montane plant populations are experiencing novel conditions due to climate change. Furthermore, climate change is causing increased climate perturbations, such as the 2012–2016 drought in the western US, remarkable in its aridity, longevity, and warmer temperatures. This drought provided an opportunity to understand how montane populations respond to extreme perturbations, including at range limits. We resurrected seeds of the endemic annual plant Erythranthe laciniata, collected in 2008 or earlier (before the drought) and in 2014 (the height of the drought), in a common garden experiment to understand how drought influenced evolution in contemporary field conditions. The study included nine populations across the species range, including range edges. Over 2,100 replicates were sown in three common gardens at natural populations at low, central, and high elevations. We recorded phenology and flower production to estimate lifetime fitness. This experiment took place in 2021, a year with low precipitation and high temperatures. We found higher fitness in the drought generation at the high garden, while both generations showed similar fitness at the central and low gardens. We detected climate adaptation at the low and high gardens, and rapidly evolved faster phenology at the high garden. Lifetime fitness was substantially lower at lower gardens overall, even for low-elevation populations. Low-elevation populations outperformed central populations at the central garden, suggesting adaptive mismatch. Together, these results indicate rapid contemporary adaptation that is beneficial at the leading edge of the species range. Nevertheless, low fitness at lower elevations may foreshadow range contraction under continued climate change.

## Introduction

Climate change is resulting in increased frequency of severe climate events like prolonged droughts, while also increasing temperatures in the growing season for many plants (IPCC 2014; Pörtner et al., 2022). Plants must adapt to such changing climates or shift their ranges to avoid extirpation and extinction (Aitken et al., 2008). Droughts are becoming more common globally (Seneviratne et al., 2021) and in Mediterranean-climate biodiversity hotspots, such as California, the focus of this study, where droughts are expected to increase in the future (Solomon et al., 2009; Essa et al., 2023). Drought is a major selective force on plant populations (Chaves et al., 2003), and influences the evolution of plant morphology, phenology, and physiology. Drought events can drastically increase mortality (Breshears et al., 2021; Marchin et al., 2022; Senf et al., 2020), which can have a long-term effect on community composition and result in range shifts or contractions (Kelly and Goulden, 2008). In order to weather severe drought events, plants need sufficient genetic variation to adapt or adaptive phenotypic plasticity to cope (Connor and Hartl 2004; Hoffmann et al., 2017).

Range limits theory suggests that adaptive responses at the cold (leading) and warm (rear) climate edges of species ranges may vary due to potential differences in abundance, isolation, or historical effects (Hampe and Petit, 2005; Pironon et al., 2017; Pennington et al., 2021; Shay et al., 2026; Sheth et al., 2026). Plants at the leading edge of a range may be able to colonize beyond-range areas that were previously uninhabitable, whereas rear edge populations may face extirpation unless adaptation to warmer, drier conditions rapidly evolves (Anderson and Wadgymar, 2020; Hampe and Petit, 2005; Mamantov et al., 2021). In montane environments, plant distributions are limited by how far their ranges can shift upward. Further, montane plants may have difficulty tracking climate change (Alexander et al., 2018) due to limited dispersal. As conditions warm, there may be niche mismatch, especially for alpine species, potentially resulting in range shifts and local extirpation (Parmesan, 2006; Parmesan and Yohe, 2003). In a comprehensive study of plant, invertebrate, and vertebrate species’ ranges, Rumpf et al. (2019) found that low-elevation populations experience extirpation at the same rate as high-elevation colonization of newly habitable areas. However, a separate analysis showed that for some species, upslope migration is outpacing new colonization—that is, that rear edges are contracting faster than leading edges are expanding (Mamantov et al., 2021). Adaptation may hold ranges stable for a time, but climate change may eventually outpace species’ ability to adapt (Jump and Peñuelas, 2005).

Rapid adaptation can mitigate rapid climate change (Anderson, 2016; Hoffmann and Sgrò, 2011; van Heerwaarden and Sgrò, 2014; Visser, 2008), and resurrection studies are helping us understand possibilities and limitations of this process, but more studies are needed to understand the fitness outcomes of rapid adaptation (Pennington et al., 2025). Rapid evolutionary changes can occur within just a few generations (Franks et al., 2018, 2007; Grant and Grant, 2002), including change in response to strong climate perturbations (Dickman et al., 2019; Gómez et al., 2018; Spurlin and Lambrecht, 2024). These rapid changes in a short period have the potential to impact the evolutionary trajectory of plant populations and may impact range dynamics such as contraction or expansion. However, rapid evolution is not always synonymous with adaptive responses; in some cases, it may lead to maladaptive outcomes, potentially reducing fitness in response to environmental stress (Battlay et al., 2023; Fussmann and Kopp, 2023). The resurrection approach provides a direct way to observe contemporary evolution, and can be further leveraged to determine whether that change is adaptive (Franks et al., 2018, 2007; Pennington et al., 2025, *in review*). Resurrection studies have thus far detected both rapid evolution and rapid adaptation in a wide variety of plants (e.g. Branch et al., 2024; Cheptou et al., 2022; Christie et al., 2023; Lambrecht et al., 2020), though remain rare in range-wide contexts (but see Anstett et al., 2021; Dickman et al., 2019; Kooyers et al., 2021) and especially in field conditions (but see Karitter et al., 2024; Sheth et al., 2026).

Changes in plant phenology are among the clearest effects of climate change impacts on plant species (Inouye, 2022; Parmesan and Yohe, 2003). Many plant species are already experiencing a change in the timing of their phenology to avoid unfavorable conditions (Walther et al., 2002), and earlier timing in phenology in particular is associated with drought escape and increased fitness in drought conditions (Farooq et al., 2009). However, although adaptive, evolutionary changes in phenology have been observed at species’ low-elevation rear edges in some cases (e.g., Perrier et al., 2025), high-elevation populations may not harbor the same adaptability in phenology as at other parts of the species’ range (Zettlemoyer and Peterson, 2021), which may hinder range expansion.

The 2012-2016 drought in the western United States provided a natural experiment of rapid evolution to climate change. Specifically studying the impact of climate change across a broad climatic gradient such as the California Sierra Nevada provides opportunities to study adaptation and strong climate impacts across species ranges (Tito et al., 2020). The drought was extreme in its strength and duration (AghaKouchak et al., 2014; Griffin and Anchukaitis, 2014); in the mountains, the lack of snowpack was exceptional (Belmecheri et al., 2016; Mote et al., 2016). More than 129 million trees died during the drought (USDA 2017). However, how this drought impacted herbaceous plants in the Sierra Nevada—the majority of the biodiversity—is still largely unknown. Plants in the Sierra Nevada are already responding to climate change through range shifts and adaptation. For example, conifers in the Sierra Nevada are responding by moving upslope to escape environments that are rapidly becoming unfavorable (Wright et al., 2016). Also, in the cutleaf monkeyflower, *Erythranthe laciniata* (formerly *Mimulus laciniatus*), changes in phenology have been observed that align with drought escape strategies (Dickman et al., 2019; Pennington and Sexton, unpublished data).

Prior resurrection research on *E. laciniata* demonstrated earlier emergence in drought generation plants, sustained after two generations, with variation across the range (Dickman et al., 2019; Pennington and Sexton, unpublished data). Flowering was found to be delayed in the first generation, but was earlier in the second (Dickman et al., 2019). This sustained change in phenology suggests that drought-adapted genotypes were selected for during drought. Further, drought-generation populations, especially in higher elevations, experienced a reduction in trait variation after drought. This is notable because *E. laciniata* has shown evidence of elevation-based adaptation (Dickman et al., 2019; Love and Ferris, 2025; Sexton et al., 2011; Shay et al., 2026), and it is possible that drought selection has reduced locally adapted genetic variation in these populations.

To better understand the fitness consequences of evolutionary responses to drought in *E. laciniata* detected in prior studies (Dickman et al., 2019; Pennington and Sexton, unpublished data), we performed a resurrection study in field conditions at multiple locations using an elevation-based, range limits context. This experiment took place in a relatively warm, dry year, providing an opportunity to test how trait evolution that occurred during the historic drought may translate to fitness variation under field conditions across the elevation gradient. We grew seeds collected before the 2012-2016 drought and seeds collected at the height of the drought in 2014 in a common garden, range-wide, transplant design to address the following questions:

1. Does the descendant generation exhibit fitness differences and increased fitness in drier, warmer conditions?
2. Do descendants maintain signals of climate adaptation or has elevation-based climate adaptation been reduced?
3. Is the elevation-based phenology cline observed in prior studies (e.g. Sexton et al., 2011; Dickman et al., 2019) expressed in the field from low to high elevations? And, which populations, if any, express evidence for rapid, adaptive phenological evolution in the field, and in which gardens?

## Materials and methods

### Study species

*Erythranthe laciniata* is endemic to western slopes of the Sierra Nevada, where it grows mainly on granite outcrops in mossy patches, along snowmelt seeps and other water sources (Fig. 1). This plant is likely to be heavily affected by climate change: it depends on spring snowmelt for water during its growing season, and as a self-fertilizing plant with a restricted range. Prior work in this system showed limited gene flow between populations with evidence for isolation by environment, but not isolation by distance (Sexton et al., 2016, 2011). There is evidence of elevation-based local adaptation across the range (Dickman et al., 2019; Love and Ferris, 2024), which was expressed most at the leading edge, which may signal a response to climate change (Shay et al., 2026). This species is ideal for addressing evolution at range limits: its range is wholly restricted to the Sierra Nevada, which allows for sampling at both the leading and rear edges of its range. The range spans almost the entirety of the elevational range of the Sierra Nevada, from below 900 m in the foothills to over 3000 m in the subalpine region, providing strong climatic variation across the range resulting in varying selective pressures.

**Figure 1.**
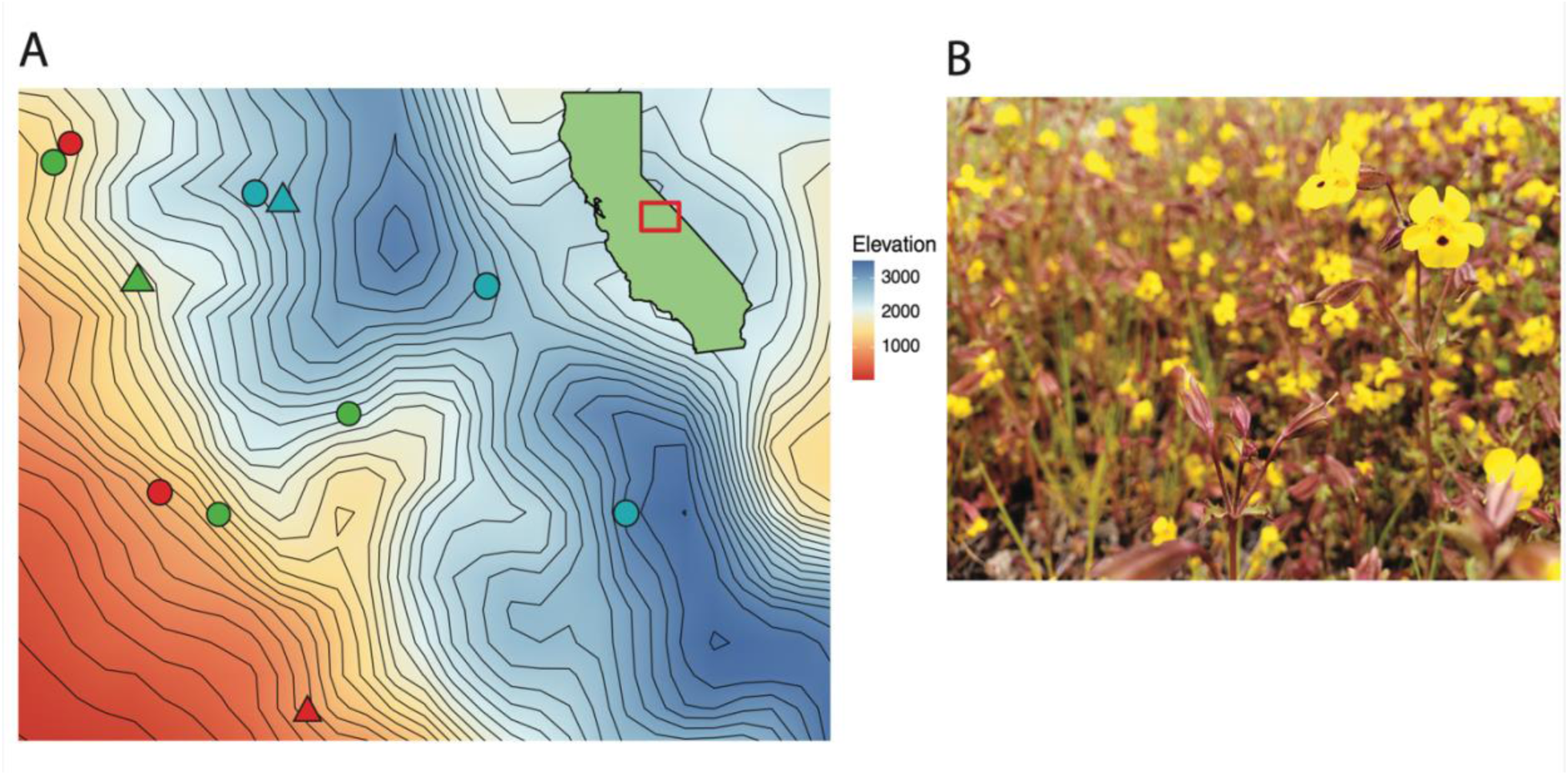
A. A contour map of the seed source populations (circles) and gardens (triangles). Low-elevation populations are colored in blue, central populations are colored in green, and high elevation populations are colored in red. Note that the low-elevation garden was also a seed-source population. Green inset map of California shows the map area outlined in red. B. *Erythranthe laciniata,* the cutleaf monkeyflower.

### Experimental Design

Seeds from the ancestral generation were collected from 2005-2008 and seeds from the drought generation were collected in 2014 (Table S1) (also, see Dickman et al., 2019). Collections from nine populations across the species’ range (Fig. 1) were refreshed for two generations in growth chamber conditions before sowing (Dickman et al., 2019). Seeds were randomly sown into small tray blocks set into the field, mimicking natural growing conditions as in Sexton et al. (2011) and Shay et al. (2026). Each block consisted of 3 rows and 6 columns into which a representative from each of the nine populations and each generation (9 populations x 2 generations) were sown; in each cell, 5 seeds were sown and later thinned to 1 plant. Sta-Green outdoor potting soil was used (Lowes, Sta-Green Potting Mix Plus Fertilizer) with a layer of sand added to the top as protective mulch. For both generations, each population was represented by 15 family lines with 3 replicates each, for a total of 810 plants at each garden and 2,430 plants in the whole experiment. Each garden had 45 trays (blocks) placed in Fall 2020 to overwinter before the growing season. Trays were placed in areas that where *E. laciniata* had grown the prior year, and were placed on water-wicking mats that keep tray soil watered while allowing root penetration into the surrounding soil. Three gardens were implemented at existing populations that represent different climate bands across the range, with gardens at 1000 m, 1555 m, and 2500 m above sea level (Fig. 1). Of the garden sites, one population, the low-elevation HWY site (Table S1) was represented by one of the nine study populations. Wildfires in 2020 prevented the use of previously planned middle and upper elevation garden sites that also represented wild, home populations in the study (Shay et al. 2026). Data were collected in Spring 2021; the 2021 water year was considered to be drier and warmer than average (CDWR 2021). We visited each garden at least four times to collect germination and phenology data, thin trays of excess plants, and to maintain the experiment over the course of the growing season. The number of reproductive branches (pedicels) was recorded at each visit, with the final count used as a fitness proxy, hereafter referred to as “total flowers” as in (Shay et al. 2026).

### Data analysis

Our experiment involved field placement of trays at the end of Fall, which allowed seeds to overwinter and experience natural vernalization. We removed data from trays that were tampered with by animals, were lost to sliding or crushing snow, or had no germination due to lack of water at the microsite by the end of the experiment (as in Sexton et al. 2011; Shay et al. 2026).

#### Does the descendant generation exhibit increased fitness in drier, warmer conditions?

To test for adaptive responses in the drought generation, we first used a two-stage hurdle model to understand survival and flower production of plants that survived to flowering. In the hurdle model, we modeled survival and flower production separately as functions of generation, source population elevation, and garden using the glmmTMB package (McGillycuddy et al., 2025) in R (R Core Team, 2025). Initially, we tested a three-way interaction (generation × elevation × garden) for each and included only significant interactions in the final models. Thus, our final model for survival included the interaction of elevation (covariate) and garden, and the interaction between elevation and generation (elevation × garden + elevation × generation), and we specified seed source population and tray nested in garden as random effects, and used a binomial family with a “logit” link. For the conditional flower count component of the hurdle model, we included only data from plants that flowered. We found a modest, significant effect of the interaction between generation and elevation and so our final model was (elevation × generation + garden), and we used a truncated negative binomial (truncated_nbinom1) distribution family and again specified seed source population and tray nested in garden as random effects.

We additionally modeled “lifetime fitness” as an integrated survival and flower count model—that is, including plants that did not survive to flowering. For this model we specified a negative binomial (“nbinom2”) distribution family and modeled the interaction of elevation and garden and the interaction of elevation and generation (elevation × garden + elevation × generation), and specified seed source population and tray nested in garden as random effects. We initially used a zero-inflated model for flower production due to a large number of zeroes in these data, but we found that zero inflation did not improve model fit and did not change significant effects. To determine the significance of factors, we used the Anova function in the car package (Fox and Weisberg 2019). We used the ggpredict function in the package ggeffects to generate figures based on our models. We predicted that descendants (drought generation), having been exposed to prolonged drought, would have higher fitness in all gardens in the relatively dry, warm climate year of the experiment.

#### Do descendants maintain signals of climate adaptation?

To allow contrasts between elevation bands, we sorted populations into elevational groups: low, central, and high, with three populations in each group. We ran a separate model with elevation group included as a covariate rather than population elevation as used in the analysis above. For this analysis we were primarily interested in how the generation of each elevation group at each garden impacts survival and lifetime fitness; thus, our final model included the three-way interaction (generation × elevation group × garden). For survival, our model used the binomial distribution with a logit link. For lifetime fitness, our model used a negative binomial distribution (“nbinom1”). Both included tray nested in garden as a random effect. The inclusion of seed source population as a random effect worsened model fit, and so we did not include it in our final model for climate adaptation. We used the Anova function in the car package (Fox and Weisberg 2019) to determine the significance of factors.

We used these models to estimate marginal means using the emmeans package (Lenth 2025) in R. We then used the contrast function in emmeans to specify a local climate group vs all others for each garden as post-hoc tests to compare generations, gardens, and elevation groups. For the low garden (1000m) contrasts, the low-elevation group was contrasted with the central- and high-elevation groups at the low garden, for the central garden (1555m), the central-elevation group was contrasted with the low- and high-elevation groups. For the high garden (2500m) contrasts, the high elevation group was contrasted with the low and central elevation groups at the high garden. Significant results suggested a difference between survival or lifetime fitness of the focal group as compared with the other groups combined (e.g., central group compared to low + high groups). Finally, we generated figures to add context to the contrast results, using the estimated marginal means. Higher survival or lifetime fitness in the focal group as compared to the other groups, and a significant contrast result, indicated broad, elevation-based climate adaptation. We predicted that ancestors (pre-drought generation) would show stronger local adaptation to the elevation gradient as compared with descendants (drought generation), as the ancestors likely had more genetic variation prior to the 2012-2016 drought, whereas the descendants likely lost genetic variation due to harsh selection imposed by the drought and therefore are less likely to show an adaptive cline to elevation.

Is the elevation-based phenology cline observed in prior studies expressed in the field from low to high elevations? And, which populations, if any, express evidence for rapid, adaptive phenological evolution in the field, and in which gardens?

Due to season-related differences among gardens in accessibility (i.e., snow cover), each garden was visited four times but on different dates. As such, we analyzed each garden separately in our phenology analysis. We first chose which visit had the highest variation in phenology responses to capture differences among populations and generations. For the low garden, this was the third visit, on April 7^th^. For the central and high gardens, it was the second visit, occurring April 21^st^ at the central garden and June 22^nd^ at the high garden. We scored phenology based on the most advanced reproductive stage observed on a plant: 0 – unemerged; 1 – sprout, lacking true leaves; 2 – vegetative, having true leaves; 3 – flowering; 4 – fruiting. As our scoring has a natural progression, we fit an ordinal logistic regression model using the polr function in the MASS package (Venables and Ripley, 2002) in R. For each model within garden, we ordered the phenological scores as factors in ascending order, and then regressed the phenological score against generation and elevation group, with block as a random effect. We generated p-values for specific effects using the Anova function in the car package. We used the predict function to estimate the probability of each phenological stage for generation and group for visualization. Based on our prior research (Dickman et al. 2019), we predicted that low-edge plants would have faster phenology than high-edge plants regardless of garden and that the drought generation would have faster phenology than the pre-drought generation regardless of garden.

## Results

### General summary and climate overview

Overall survival was 21% across the experiment, but was higher at the high garden (38.3%) (Table 1). Flower production was also higher at the high garden (Table 1).

**Table 1.**
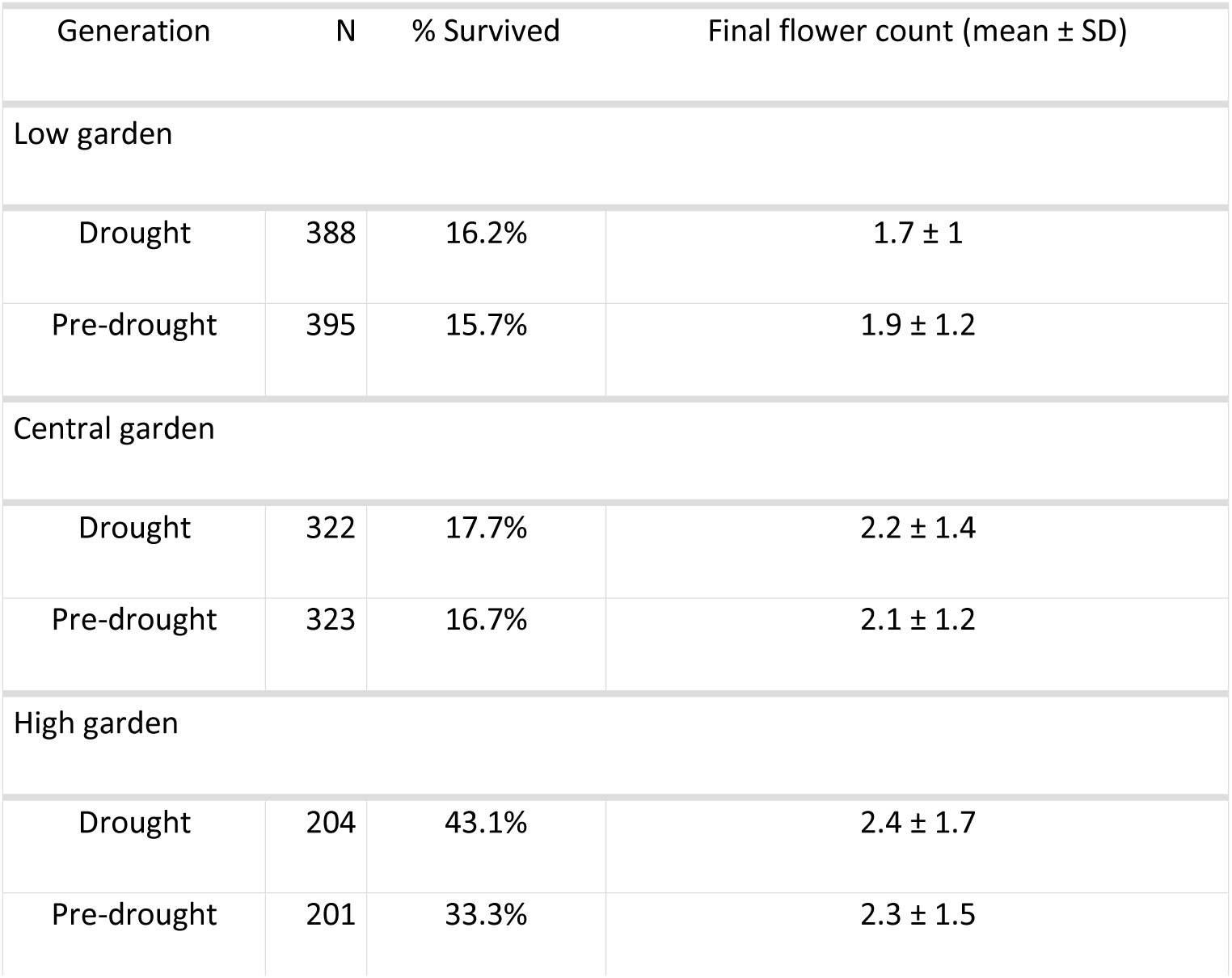
Summary of sample sizes, survival rates, and average final flower counts at each garden for each generation.

The gardens were sown in Fall 2020 and visits began in early 2021, meaning the experiment took place in water year 2021 (October 1, 2020 – September 30, 2021). Water year 2021 was notably dry and warm, and at the time was the second driest year on record and the second warmest year on record for statewide temperature (CDWR 2021). Temperatures were warmer in 2021 than in either collection years for this experiment, especially at the high garden, though temperatures were warmer than the 25-year average at all time-points (Fig. S1). Precipitation was also lower for almost all time-points, except for a few source populations in the pre-drought collection year (Fig. S2).

#### Does the descendant generation exhibit increased fitness in drier, warmer conditions?

##### Survival

We found that although most factors affected survival significantly, garden had the strongest effect (Fig. 2). Garden (p = < 0.001) and the interaction between garden and elevation (p = < 0.001) were significant, which was largely driven by higher survival in the high garden. The results for elevation (p = 0.002), generation (p = 0.12, not significant), and their interaction (elevation × generation: p = 0.05), show that the adaptive cline shifts by generation, driven by the high garden. This aligns with our hypothesis of adaptive evolution during the drought. The highest elevation populations consistently showed higher survival probabilities in the drought generation than in the pre-drought counterpart (Fig. 2). Overall, survival decreased as source population elevation increased at the low (1000m) and central (1555m) gardens, and for the pre-drought generation in the high garden. Survival was highest for all populations at the high elevation garden (2500m), and this was especially pronounced in the two highest elevation source populations (Fig. 2). These patterns indicate an adaptive elevation survival cline as observed in Shay et al. (2026). Further, the regression lines of fitness versus elevation for both generations had a negative slope, illustrating that lower elevation genotypes perform better at all three gardens regardless of generation—with the exception of the drought generation at the high garden, which had a positive regression slope, providing evidence of rapid, adaptive evolution being expressed in the field in this dry, warm year.

**Fig. 2.**
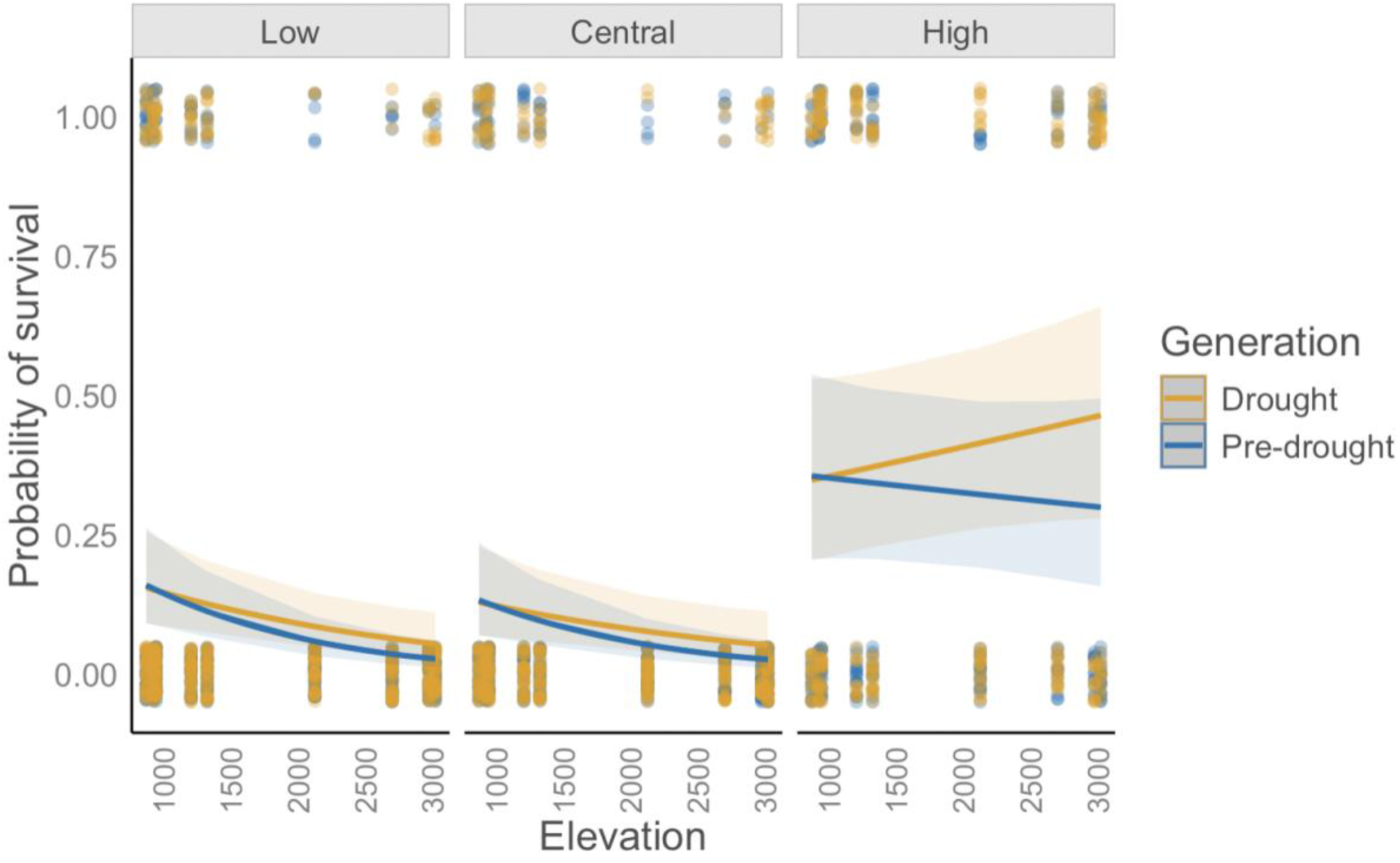
Probability of survival predicted from binomial model. Elevation is on the x-axis with survival probability on the y-axis. The figure is faceted by garden, with the lowest elevation garden (1000m) on the left, the central elevation garden (1555m) in the center, and the highest elevation garden (2500m) on the right. The two generations are separated by color with the pre-drought/ancestor generation in blue and the drought/descendant generation in orange. Shaded areas surrounding prediction lines are the 95% confidence intervals.

##### Flower production

###### Conditional flower production

When modeling flower production of only plants that survived to flowering in the hurdle model, we did not find significant effects from our predictors (Table S2). The interaction between generation and elevation was weakly significant (p = 0.06). This interaction was likely driven by the drought generation of the higher elevation populations consistently producing more flowers than the predrought generation (Fig. 3), which lends limited support to our hypothesis of adaptive evolution during the great drought.

**Fig. 3.**
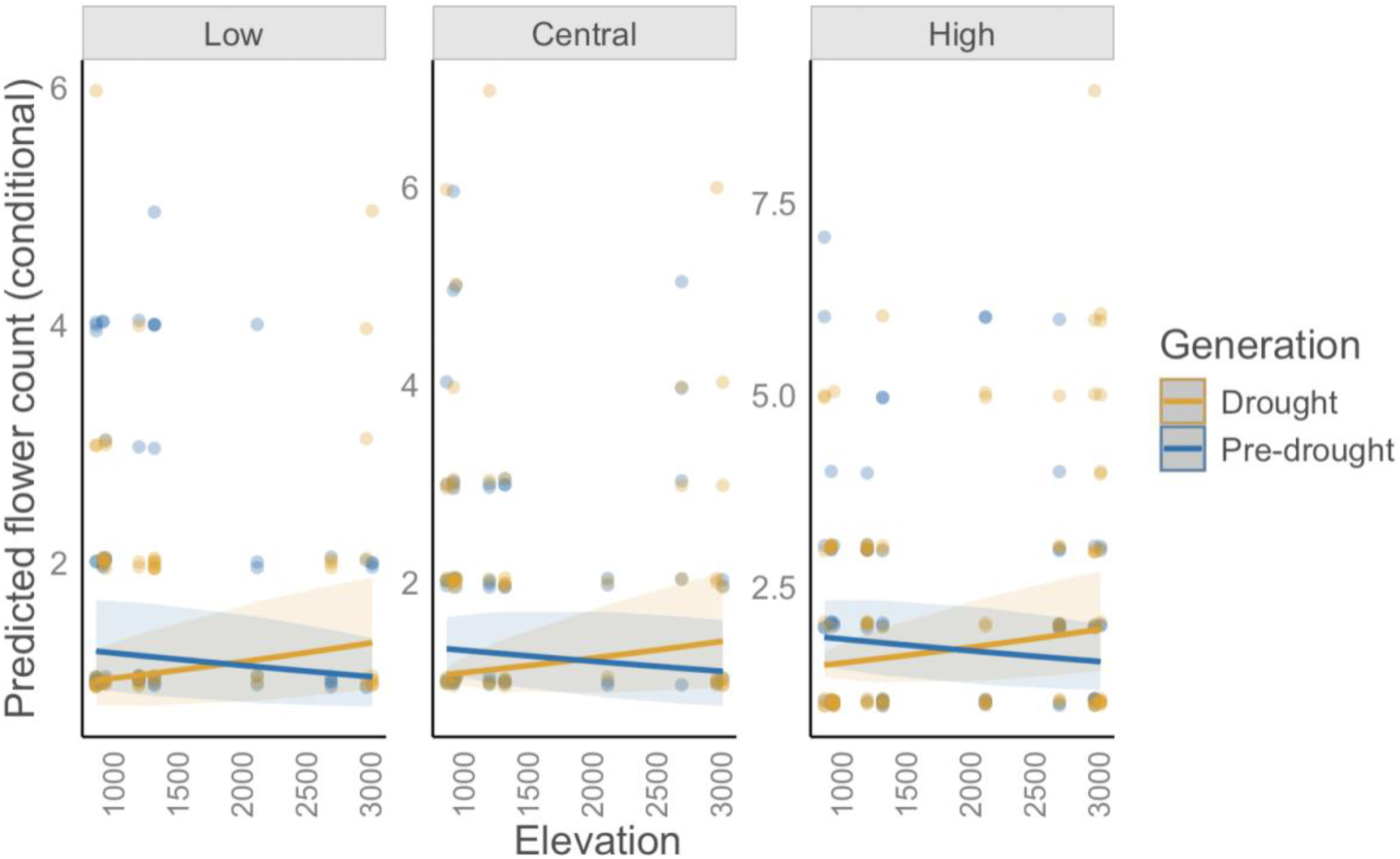
Predicted flower count from the count portion of hurdle model. Elevation is on the x-axis with survival probability on the y-axis. The figure is faceted by garden, with the lowest elevation garden (1000m) on the left, the central elevation garden (1555m) in the center, and the highest elevation garden (2500m) on the right. The two generations are separated by color with the pre-drought/ancestor generation in blue and the drought/descendant generation in orange. Shaded areas surrounding prediction lines are the 95% confidence intervals.

###### Integrated lifetime fitness

When modeling lifetime fitness, which included survival and flowering into one measure, we found stronger patterns from our predictors. Lifetime fitness did not vary significantly by generation (p = 0.57), but was significantly affected by garden (p = < 0.001), elevation (p = 0.02) and their interaction (generation × elevation, p = 0.02), and this relationship can be seen again in the positive regression slope representing higher flower production in the drought generation than the pre-drought generation at the high garden (Fig. 4). The interaction between elevation and garden was also significant (elevation × garden, p = <0.001), which can again be seen in the much higher flower counts at the high elevation garden (Fig. 4). Overall, we saw a pattern similar to survival but with lower magnitudes of reduced flower production with increased source population elevation at the Low and Central gardens. At the high-elevation garden, lifetime fitness was an order of magnitude higher than the other two gardens (Fig. 4). Finally, we again saw a negative regression slope in the pre-drought and drought generations in the low and central gardens, but a positive slope for the drought generation at the high garden, further supporting our findings of rapid adaptation to warmer, drier conditions.

**Fig. 4.**
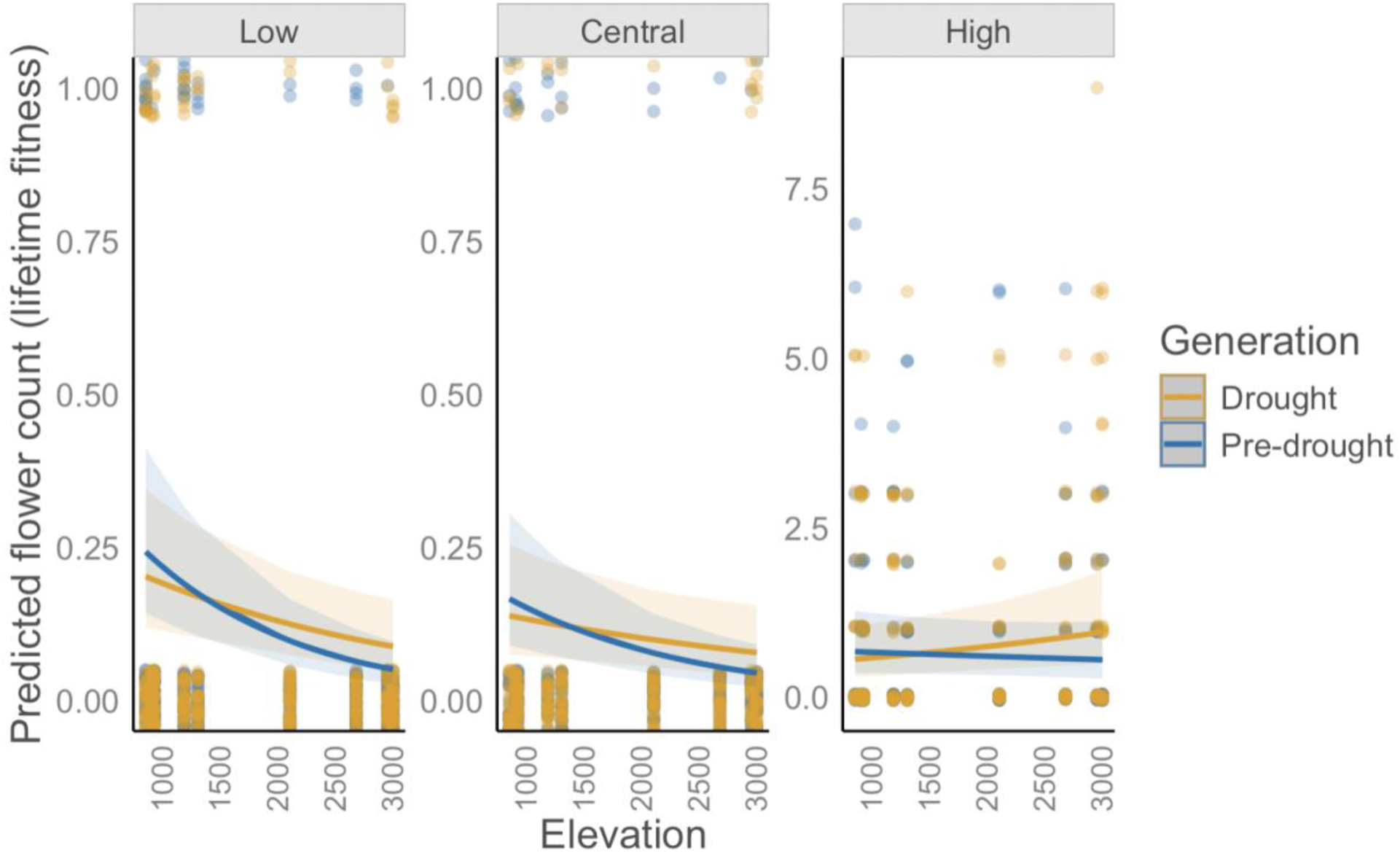
Predicted flower count from the lifetime fitness model. Elevation is on the x-axis with survival probability on the y-axis. The figure is faceted by garden, with the lowest elevation garden (1000m) on the left, the central elevation garden (1555m) in the center, and the highest elevation garden (2500m) on the right. The two generations are separated by color with the pre-drought/ancestor generation in blue and the drought/descendant generation in orange. Shaded areas surrounding prediction lines are the 95% confidence intervals. Note the y-axis scale difference for the low and central gardens compared to the high garden.

#### Do descendants maintain signals of climate adaptation?

##### Survival

In the survival model with elevation group as a fixed effect rather than population elevation, we found that generation (p < 0.001), elevation group (p = 0.001), and garden (p < 0.001) were all significant factors in the model, as well as the interaction between elevation group and garden (p = 0.002) (Table S3). Post-hoc contrasts showed that signals of elevation-based climate adaptation were maintained at the low garden, with both generations for the low group at the low garden having higher survival than the other groups (Table 2, Fig. 5). Broad climate adaptation in this sense expressed as survival was not detected at the central or high gardens (Table 2). Survival for the high group drought generation was higher at the high garden overall.

**Figure 5.**
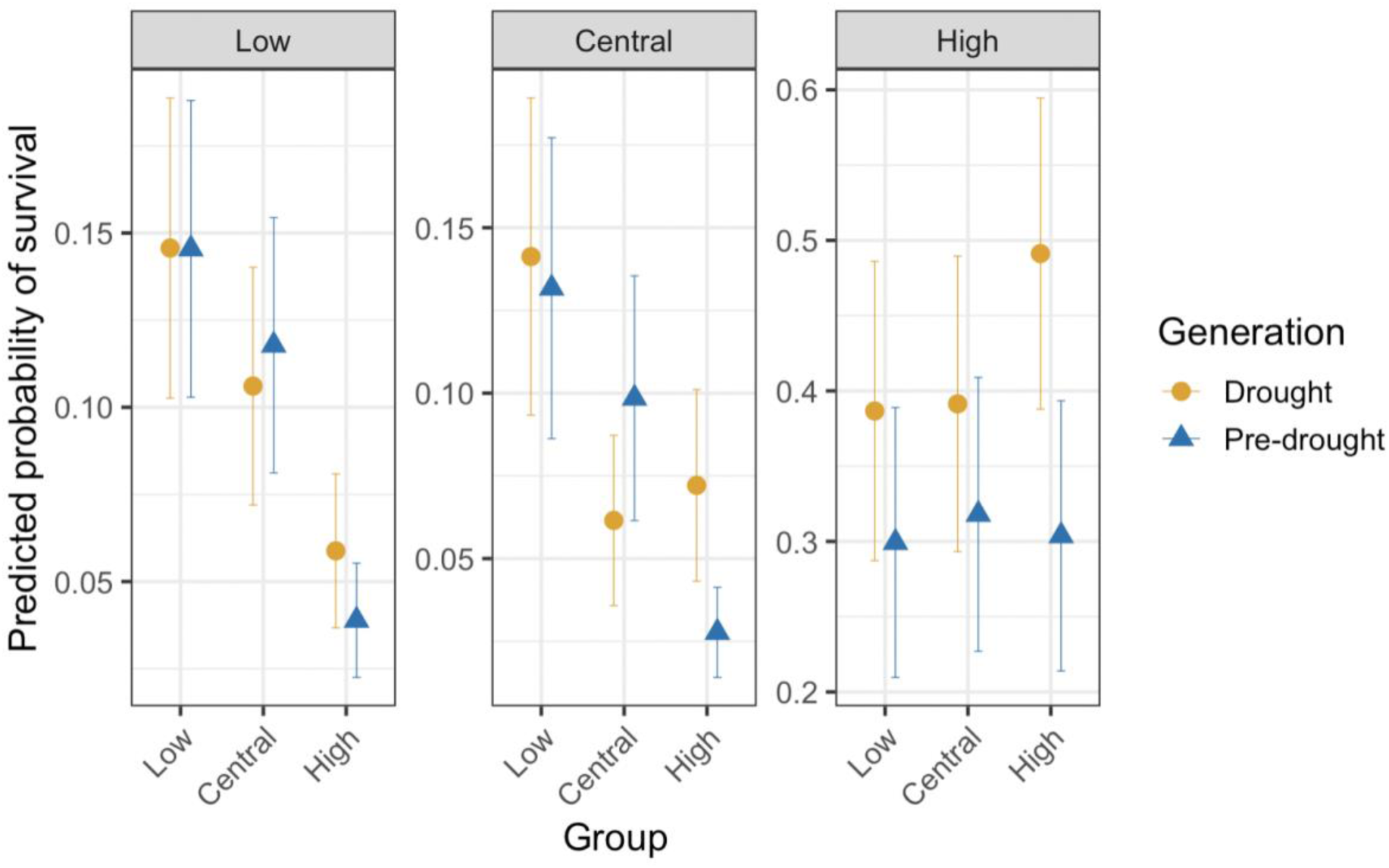
Probability of survival by garden, elevation group, and generation. Group is on the x-axis and survival probability is on the y-axis. Note scale difference for the low (1000m) and central (1555m) gardens as compared to the high garden (2500m).

**Table 2.**
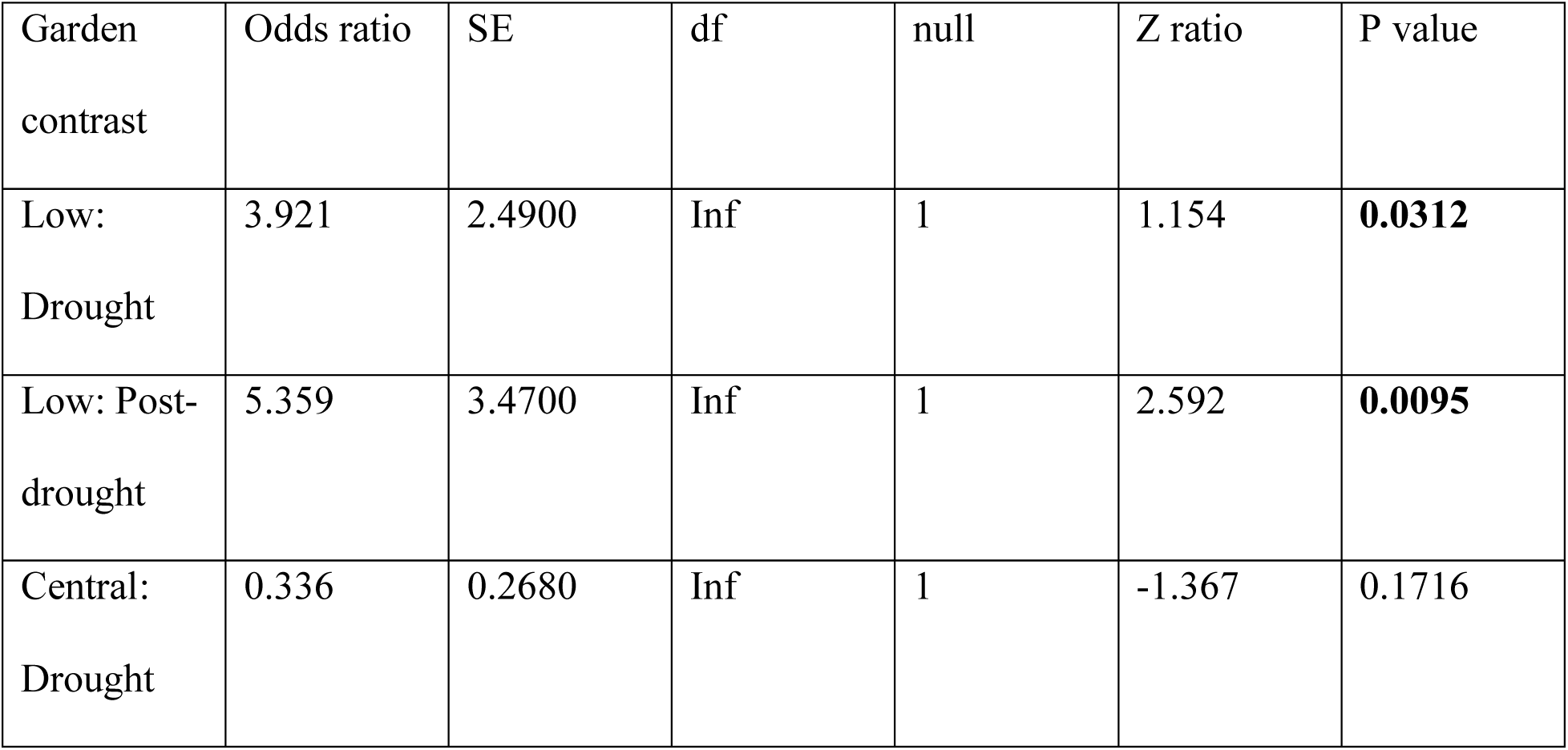
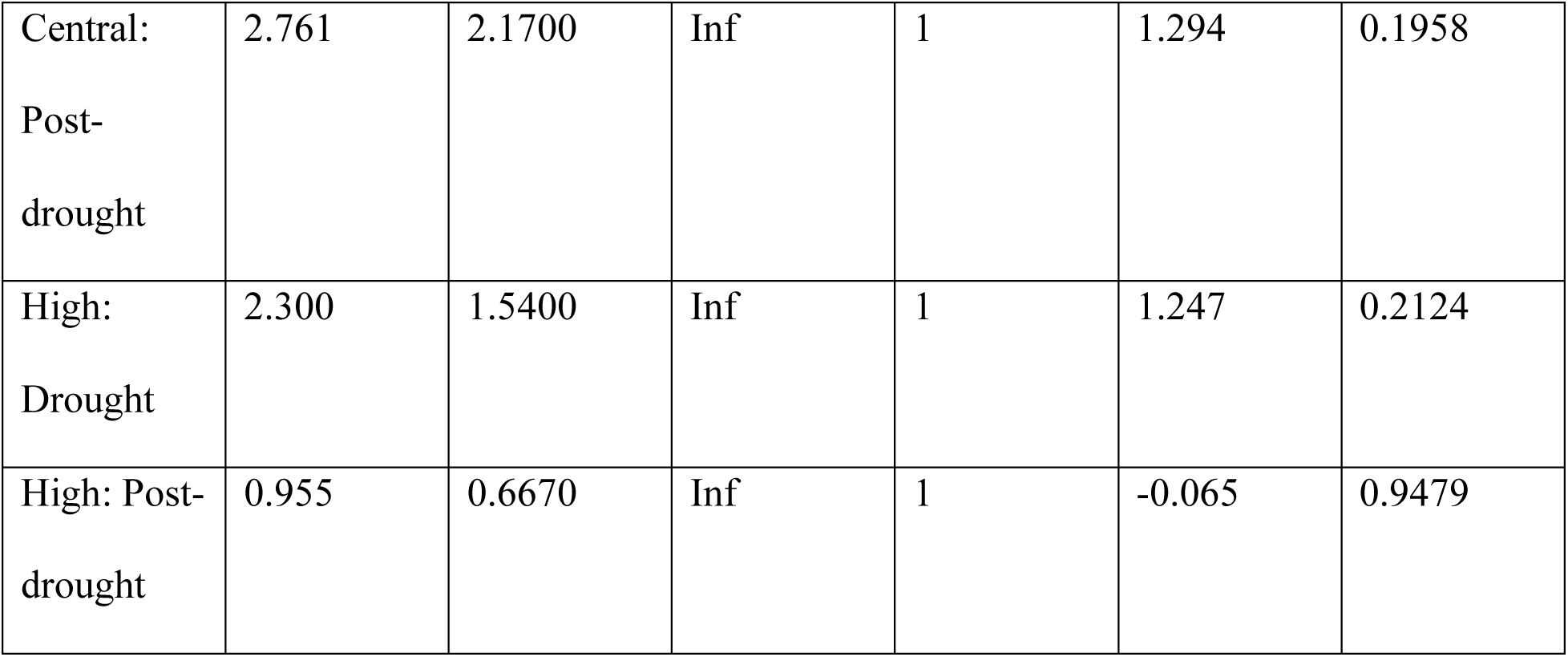
Results of post-hoc contrasts showing climate adaptation in survival. Within each garden, each generation was contrasted separately.

##### Lifetime fitness

In the lifetime fitness model with elevation group as a fixed effect rather than population elevation, we found that elevation group (p = 0.009), garden (p < 0.001), the interaction between generation and elevation group (p = 0.03) and the interaction between elevation group and garden (p < 0.001) were significant factors (Table S3). Post-hoc contrasts again showed that elevation-based climate adaptation was maintained at the low garden, without overall evidence at the high garden (Table 3). The low group in both generations had higher lifetime fitness than all other groups at the low garden (Fig. 6). Mirroring survival, flower count was higher at the high garden overall, and in the drought generation the difference between the high elevation group and all others is significant (Table 3). Thus, the high elevation group in the drought generation had greater lifetime fitness than the two other drought generation groups at that garden, suggesting that the rapid adaptation occurred during the 2012-2016 drought.

**Figure 6.**
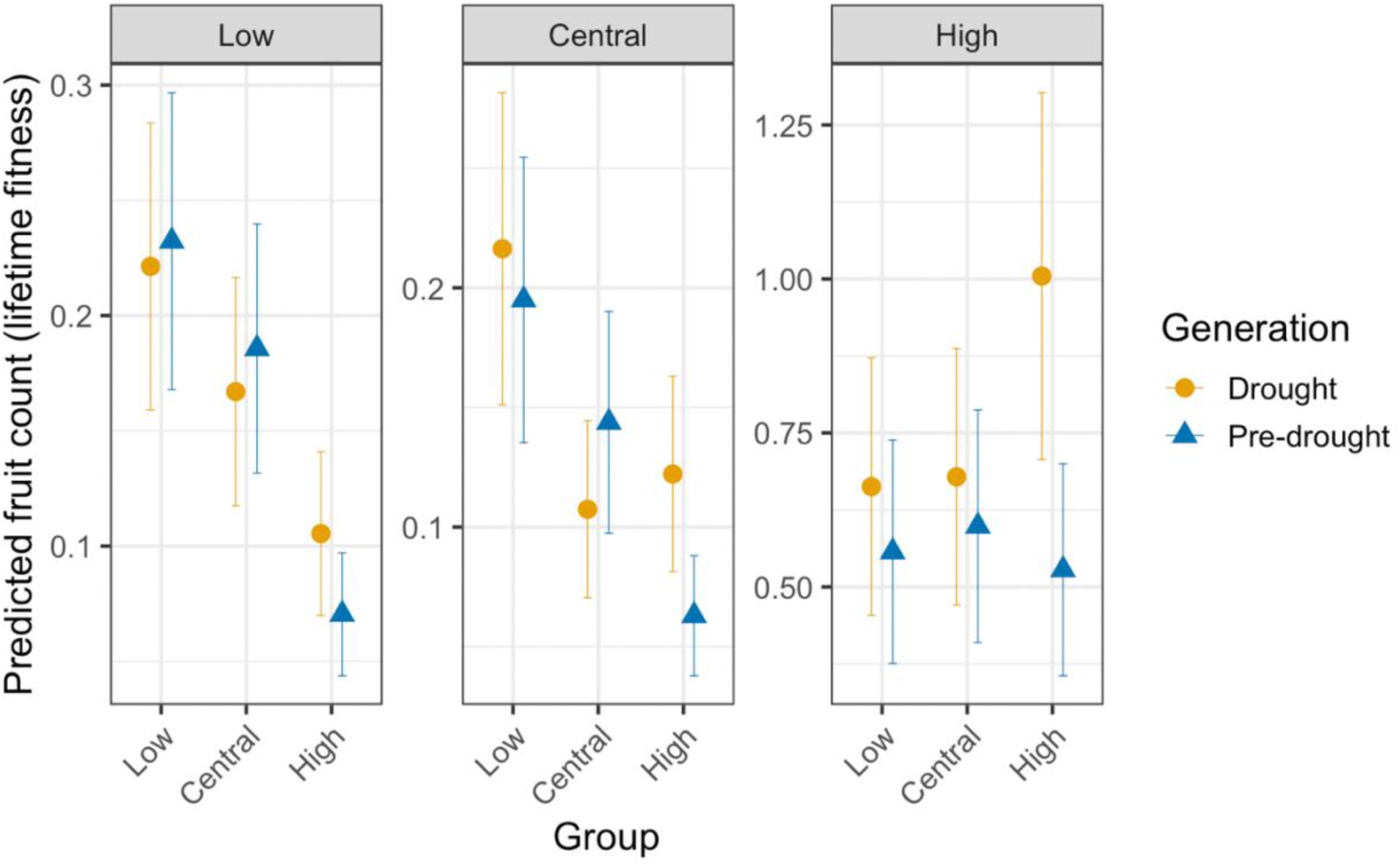
Predicted flower counts from lifetime fitness model by garden, elevation group, and generation. Group is on the x-axis and predicted flower count is on the y-axis. The drought/descendant generation is in yellow with the pre-drought/ancestor generation in blue. Note y-axis scale difference for the low (1000m) and central (1555m) garden compared to the high garden (2500m).

**Table 3.**
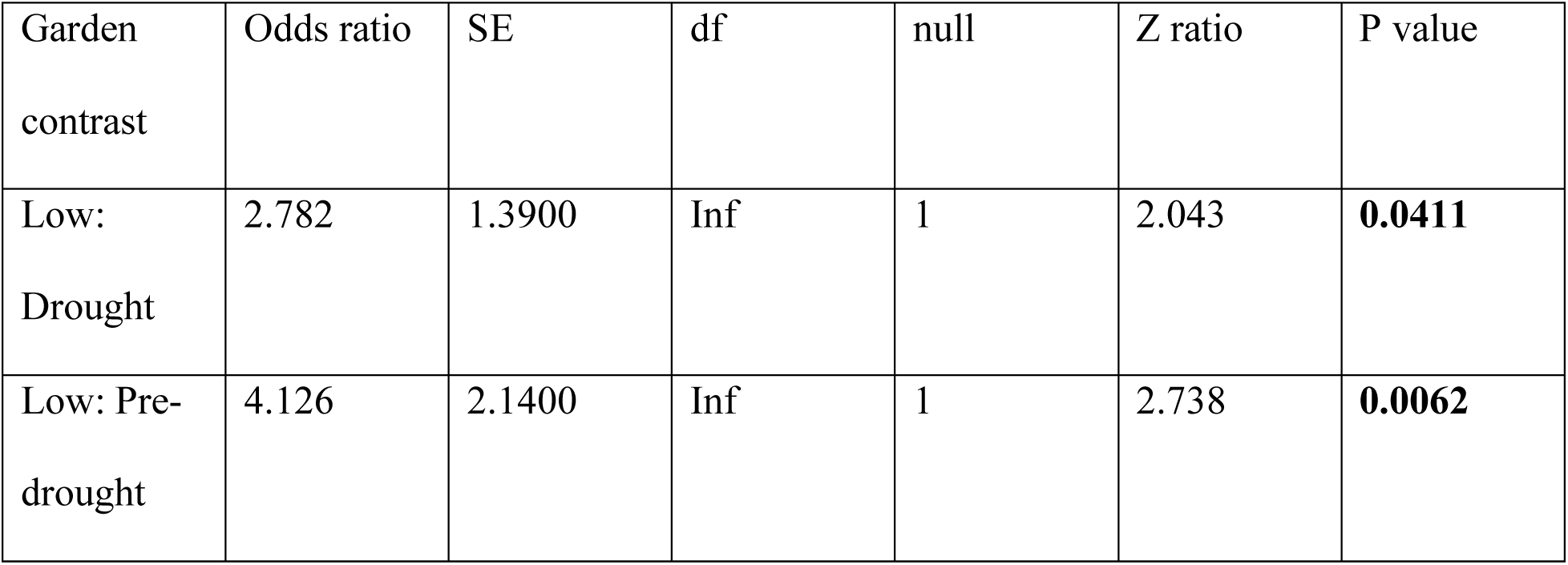
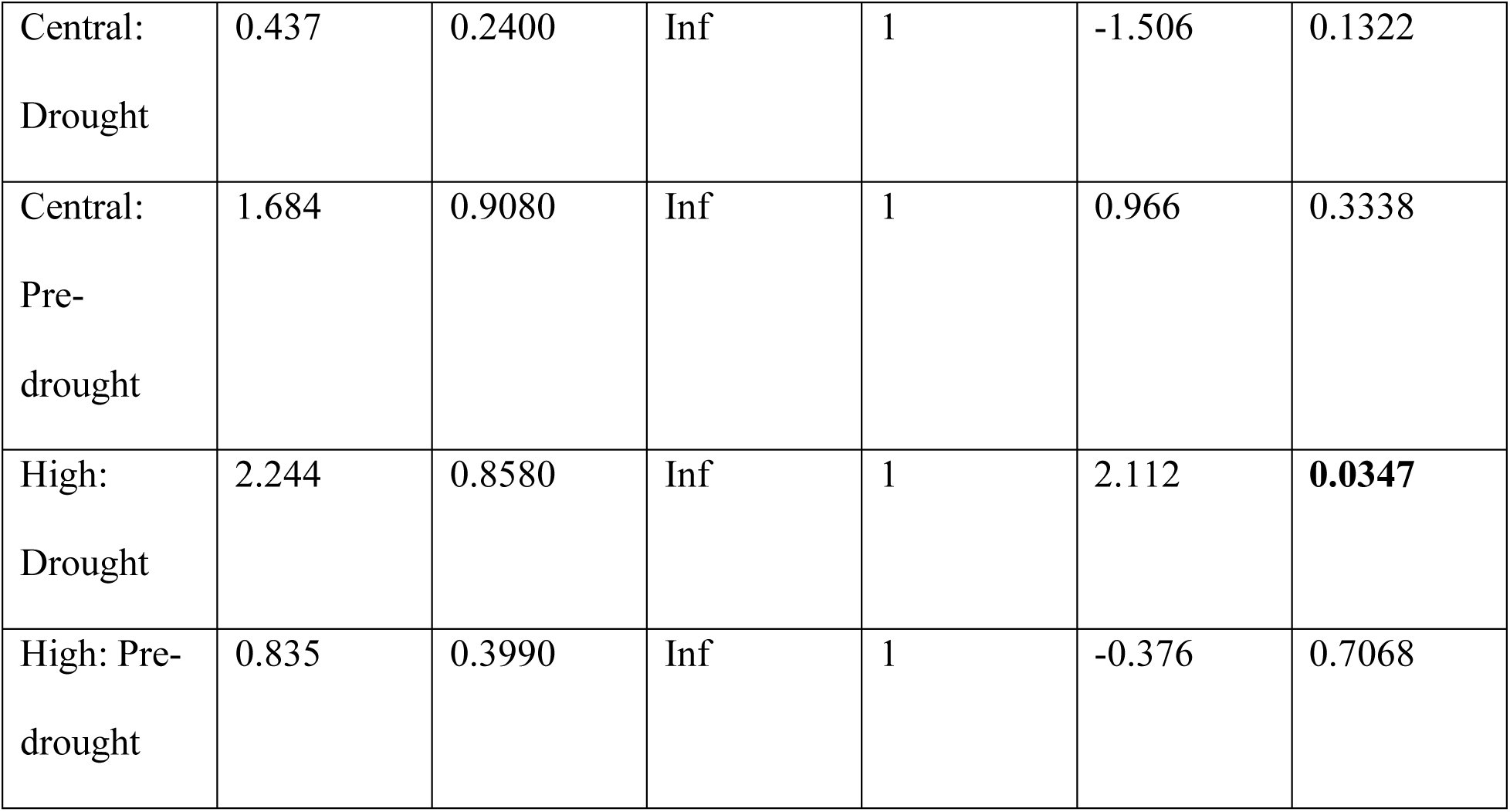
Results of post-hoc contrasts showing climate adaptation in lifetime fitness, mainly at the low-elevation garden (1000m). Within each garden, each generation was contrasted separately.

#### Is the elevation-based phenology cline observed in prior studies expressed in the field from low to high elevations? And, which populations, if any, express evidence for rapid, adaptive phenological evolution in the field, and in which gardens?

In our ordinal regression models of phenology, we found elevation group to be significant at the low (p = 0.009) and central (p < 0.001) gardens, whereas generation was not significant (Table S4). At the high garden, elevation group was not significant, whereas generation was significant (p = 0.04) (Table S4). At the high garden, the drought generation had more advanced phenology than the pre-drought generation (Fig. 7). At the central and low gardens, the low and central groups had more advanced phenology than the high elevation group, regardless of generation, indicating phenology as a trait associated with elevation-based climate adaptation.

**Figure 7.**
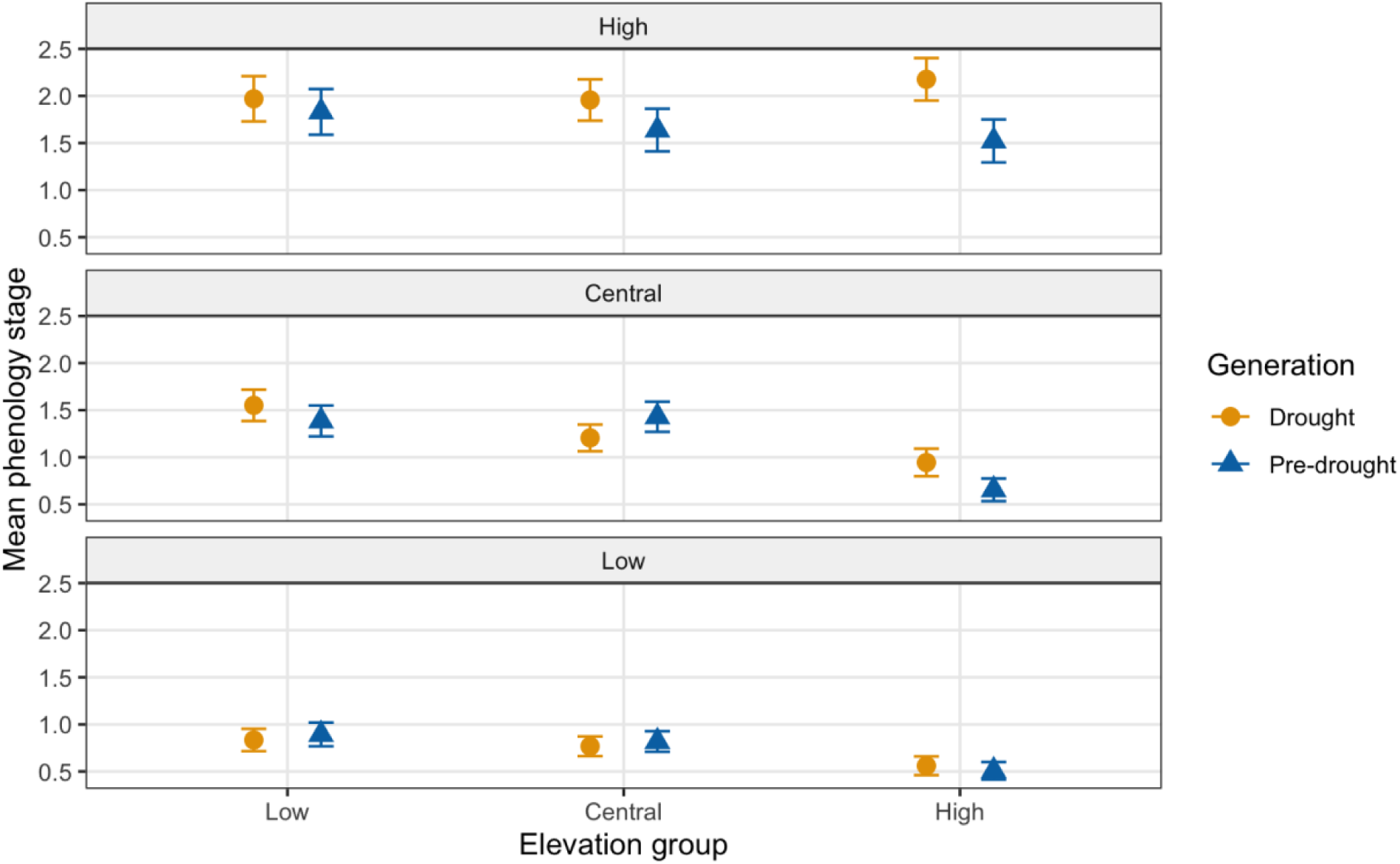
Mean phenology of each garden during sampling periods when phenological variation was greatest among plants within each garden. Phenology stage was measured as 0, unemerged; 1, sprout; 2, vegetative; 3, flowering; 4, fruiting. Bars represent standard error. The drought generation is indicated by circles, and the pre-drought generation is indicated by triangles.

## Discussion

In this study, we aimed to understand how rapid evolutionary change observed during a historic drought affected the survival and fitness outcomes during a warmer, drier field year in a native, annual forb. We implemented a resurrection study in a common garden framework across the elevational range of our focal species, *E. laciniata*. We found the most pronounced adaptive response in high-elevation populations when grown at the highest garden, which exhibited higher survival, lifetime fitness, and advanced phenology. We found that climate adaptation is maintained in the low garden across generations, and that in this dry, warm experimental year, lower populations had relatively high fitness at central and high elevations, consistent with adaptive lags during climate change. Nevertheless, high-elevation populations of the drought generation exhibited the highest fitness at the high garden and have likely rapidly evolved to maintain climate adaptation at higher elevations. To our knowledge, this study is the first time rapid, adaptive evolution has been demonstrated at the leading edge of a species range under natural conditions.

### Does the descendant generation exhibit increased fitness in drier, warmer conditions?

We found increased survival in the descendent, drought generation, but expressed only in high-elevation populations at the high-elevation garden. Survival at the high garden was higher overall regardless of generation or source population elevation. The experiment took place during a warmer and drier-than-average year—more of which are expected as climates warm (Mallakpour et al 2018). In warm, dry conditions like the 2021 growing season, it is expected that if rapid adaptation to drought occurred during the “big drought,” the drought generation would have higher fitness (Franks et al 2007). Lifetime fitness was significantly higher in the drought generation as with survival, but again, mainly at the high garden. The high-elevation garden produced more flowers overall, regardless of elevation group, suggesting that more favorable conditions for this species occur at higher elevations; this finding may indicate global warming responses (i.e., population increases at the leading edge, and decreasing at the rear edge), and is similar to prior findings in *E. laciniata* (Sexton and Dickman, 2016; Shay et al., 2026). In a warm, dry year these patterns are not surprising—however, if populations have sustained low recruitment when exposed to harsh conditions, extirpation is possible, potentially resulting in range shifts. Moreover, the faster phenology that evolved rapidly (discussed below) across most populations examined in this study (Dickman et al. 2019; Pennington and Sexton, *unpublished data*) did not translate into higher fitness or survival in the drought generations at the lower gardens (except for high-edge populations in lower-elevation gardens, which showed higher fitness and survival in the drought generation than the pre-drought cohort, but underperformed compared to low-edge populations at the lower gardens (Fig. 6). This lack of rapid, adaptive response for local, low-edge populations in the low garden may signal limits to adaptive potential in rear-edge, and other populations generally, in the face of severe climate events.

Range shifts as a result of climate change have been documented globally (Angert et al., 2011; Bradley et al., 2024; Chen et al., 2011; Zu et al., 2025). For example, in the Santa Rosa Mountains of southern California, from 1977 to 2007 researchers saw an upward trend in species distributions—populations are contracting or disappearing in lower elevations where they used to be abundant (Kelly and Goulden, 2008). Nevertheless, in many montane environments, climate change is thus far increasing biodiversity as a result of invasion or range shifts (Dornelas et al., 2014), and may result in the formation of novel communities (Alexander et al., 2015; Williams and Jackson, 2007). For *E. laciniata*, which lives in a harsh environment, thereby escaping competition (DeMarche et al., 2013; Ferris and Willis, 2018), community changes within its habitat may not be as dramatic as in more productive habitats. For this species and others like it, understanding fitness responses to climate change will be critical to their conservation.

### Do descendants maintain signals of climate adaptation?

Shay et al. (2026) demonstrated strong climate adaptation in leading-edge populations of *E. laciniata* during the 2009 growing season in the California Sierra Nevada. In this study in 2021, which was warmer and drier than 2009, we found stronger evidence of elevation-based climate adaptation at the low-edge garden—low-elevation populations had higher survival and lifetime fitness at the low garden compared to central and high-elevation populations. Additionally, the low population group outperformed the central population group at the central garden (Figs. 5, 6), which could signal climate mismatch at central elevations due to rapid climate change. Further, we also recovered a signal of climate adaptation at the high garden in lifetime fitness in 2021, but only in the drought generation; the high-elevation drought generation had higher lifetime fitness at the high garden than the central and low elevation populations, suggesting that rapid evolution at higher elevations during the drought may have maintained or “rescued” the adaptive cline at higher elevations. However, it is important to acknowledge that the three high-elevation populations in this study come from elevations above the 2021 high garden at Tenaya Lake (TL = 2500m, versus 2774m, 3049m, and 3095m for ML, ME, and HE populations, respectively). Thus, the lack of strong climate adaptation near the leading edge observed in the pre-drought generation in this experiment, compared to Shay et al. (2026), may be due in part to high-edge plants underperforming in the high (TL) garden in 2021. In a warm, dry year, these findings signal that rapid evolution undergone during drought provided an advantage in contemporary conditions. This, coupled with upwards migration from lower-elevation populations, may help higher-elevation populations adapt as climates continue to warm.

Recent climate change may be eroding patterns of local adaptation at the warmer areas of species ranges as the climates there become more stressful over time (Anderson and Wadgymar 2020; Shay et al. 2026). Although the low-elevation populations had higher mean fitness at the high garden than when grown at their low, local elevations, both generations of the low group maintained adaptive, elevation-based patterns in the low garden—that is, the low elevation populations had higher survival and lifetime fitness at the low garden than the central and high elevation groups. This finding partially supports adaptation expectations from a meta-analysis of local adaptation in common garden and transplant studies that examined population fitness, habitat quality, spatial range position, and climatic position: Bontrager et al. (2021) reported that warm-edge populations are more likely than cold-edge populations to show adaptive patterns, but that adaptation may be constrained at range edges in general due to reduced habitat quality. In contrast to general findings that local adaptation is reduced at high-elevation populations, either due to more recent colonization or reduced genetic variation, we found strong signals of climate adaptation at the at the leading edge in *E. laciniata* (Shay et al. 2026; this study).

Resurrection studies that include the leading edge of a species range are uncommon, but other studies in *Erythranthe* have examined leading edge populations. Sheth et al. (2026) found climate maladaptation in northern populations of *E. cardinalis*, with northern populations underperforming at a northern garden during the first year of growth when compared with central and southern populations. Similarly, Anstett et al. (2021) found little evolutionary response to drought in northern populations of *E. cardinalis*, but other research in this system has since found evidence of evolutionary rescue rebounding population growth rates in populations where drought-associated alleles have increased (Anstett et al. 2025). Differences in responses at the leading edge of these two congeners may be related to any number of factors, including life history differences—*E. cardinalis* is a primarily outcrossing perennial species, whereas *E. laciniata* is a primarily self-fertilizing annual.

### Do low-edge plants show faster phenology than high-edge plants? Did faster phenology evolve over the drought in edge populations?

Although lower-elevation populations had faster phenology than high-elevation populations overall, we found evidence for rapid evolution of faster phenology in all populations when grown at the high-elevation garden where plants from all populations were able to grow much larger—this was expressed most strongly in the high-elevation populations, which exhibited the fastest phenology of all elevation groups at the high garden (Fig. 7). In a greenhouse study, Dickman et al. (2019) found earlier emergence for most of the same nine *E. laciniata* populations in this study, including populations from the three highest and three lowest elevations (i.e., rear-edge and leading-edge populations). In this study, we found that phenology was still faster in the drought generation of these edge populations, but this was only expressed at the central and high gardens (Fig. 7). Drought-generation plants had greater survival and lifetime fitness, but mainly in high-elevation populations at the high garden. Taken together, we provide evidence that phenology changes that evolved in the 2012-2016 drought may confer higher survival and lifetime fitness under drier, warmer conditions, and this rapid adaptive evolution was mainly detected at the high elevation, leading edge of this species’ range during an especially warm, dry year in the field. These findings are consistent with findings that evolutionary rescue may help populations weather intense climate events where adaptive alleles are favored (Anstett et al., 2025; Bay et al., 2018).

Phenology such as flowering time is one of the most examined traits in resurrection studies, and annual plants are generally evolving earlier flowering time in response to drought and warming events (Pennington et al., 2025). In fact, earlier flowering was found to be adaptive in the seminal plant resurrection study (Franks et al., 2007), but the directionality of phenology varies by system. For warm, rear-edge populations of *Cyanus segetum*, earlier phenology conferred higher fitness in a field context (Valencia-Montoya et al., 2021). Conversely, a greenhouse experiment revealed that slower phenology evolved across the species range of *Erythranthe cardinalis* in response to the great drought (Anstett et al., 2021). Although southern populations of *E. cardinalis* did flower later than central and northern populations in a southern garden, rapid evolution in phenology that translates to fitness differences have so far not been observed in the field (Sheth et al. 2026).

Whether evolution in phenology or phenotypic plasticity in phenology will be able to help plants keep pace with climate change to bolster population fitness and avoid extirpations is an open question (Anderson et al., 2012; Duputié et al., 2015). Further, plasticity in phenology rather than evolutionary change as a likely response to rapid climate change is hypothesized to be more common (Ramirez-Parada et al., 2024). Phenotypic plasticity in phenology has been commonly documented in resurrection studies (e.g. Anstett et al., 2021; Lambrecht et al., 2020; Rauschkolb et al., 2022), but there is a clear need for more studies of rapid evolutionary potential in the natural conditions. Here, we find evidence that rapid adaptation may be assisting some plant populations to weather modern, severe climate events caused by climate change, including at edge populations, but the evolutionary potential to help species continue “running to stand still” clearly has limits (Jump and Peñuelas, 2005).

## Supplement

**Fig. S1.**
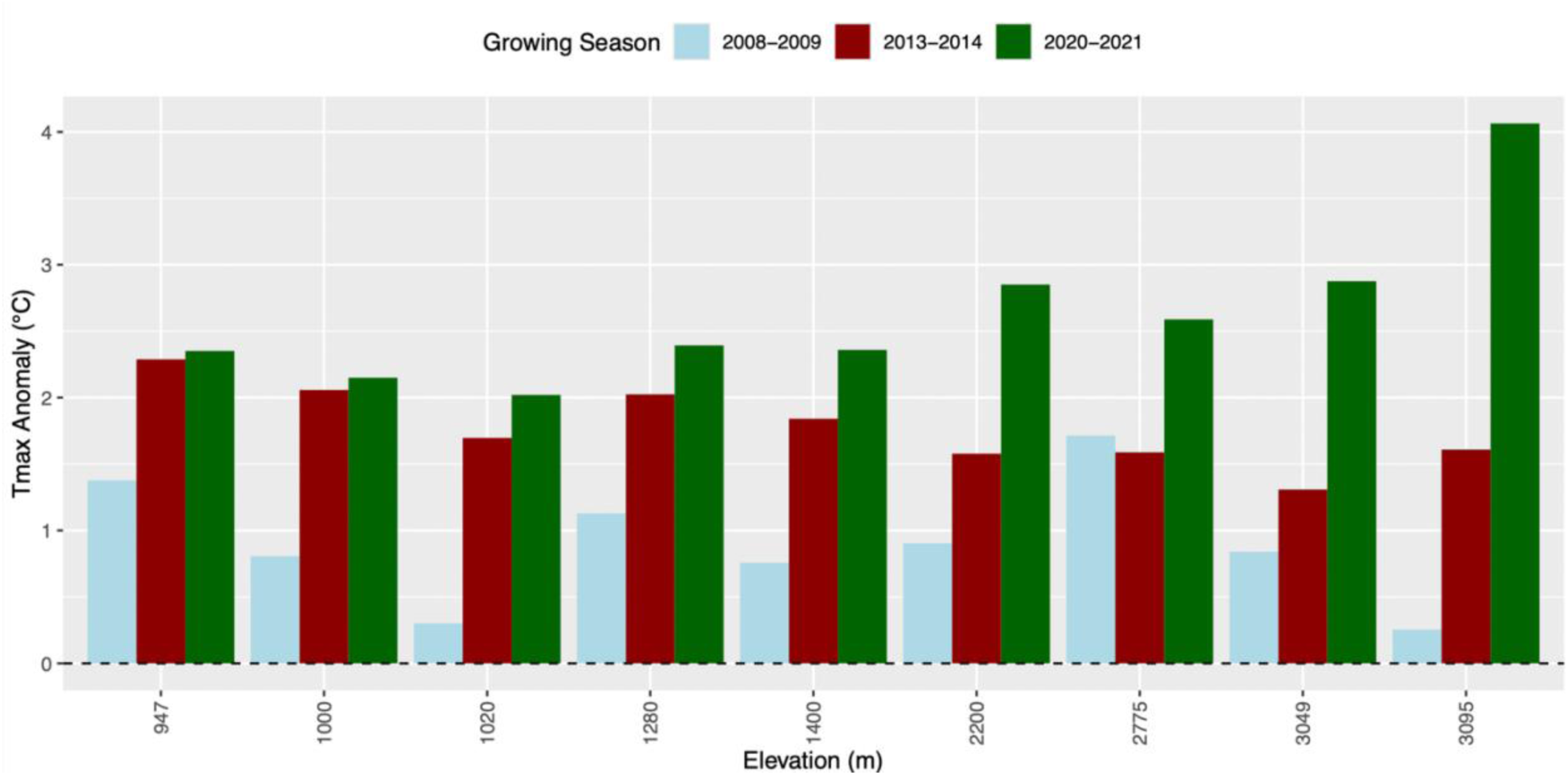
Temperature difference from 25-year average for each collection water year and the garden water year

**Fig. S2.**
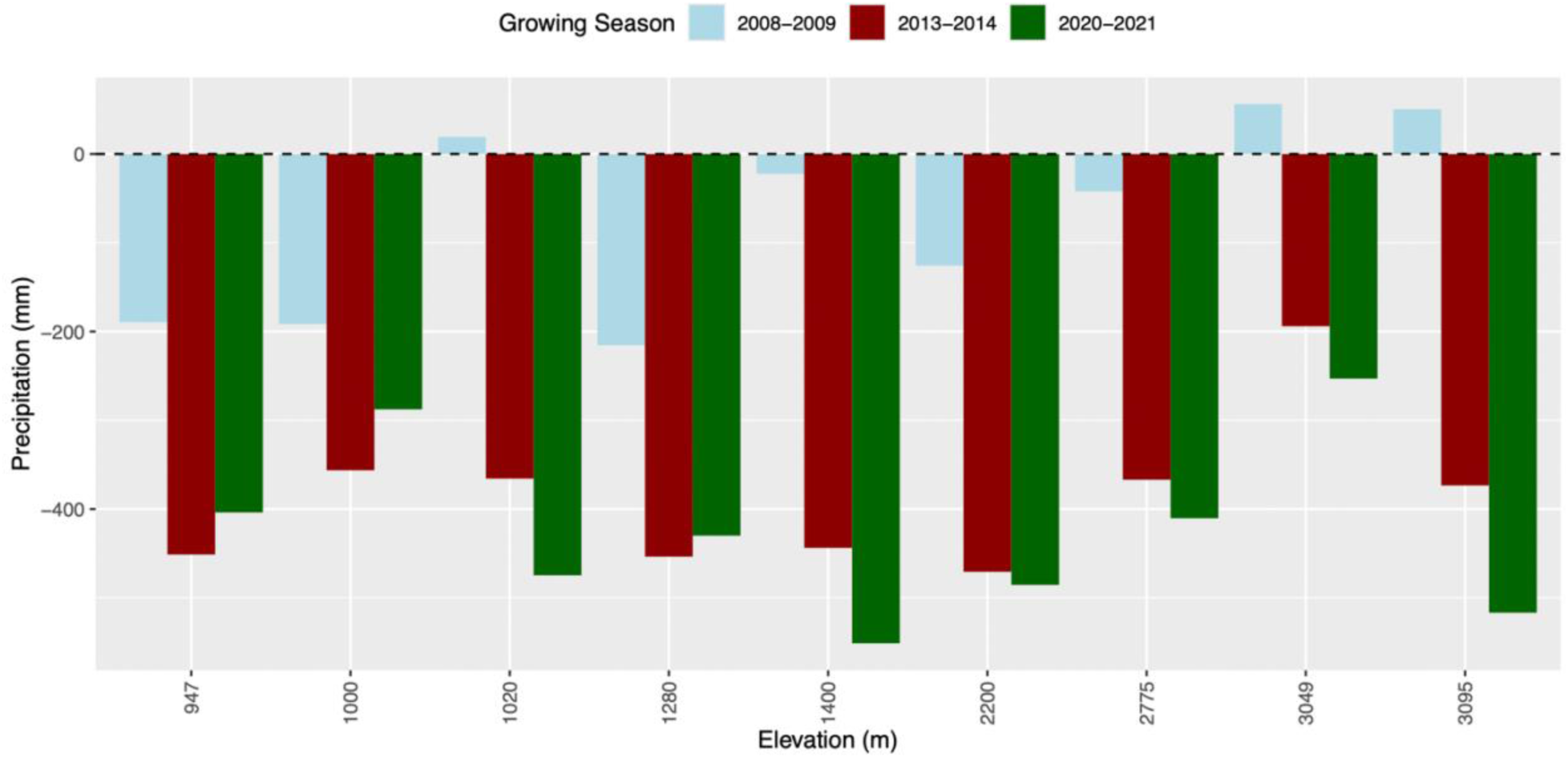
Precipitation difference from 25-year average for each collection water year and the garden water year

**Table S1.** Population information with collection dates for the pre-drought and drought generation.

| **Population Code** | **Population Name** | **Elevation (meters)** | **Pre-drought Collection Year** | **Drought Collection Year** |
| --- | --- | --- | --- | --- |
| R | R Property | 947 | 2006 | 2014 |
| HWY | Hwy 168 | 1000 | 2006 | 2014 |
| HH | Poopenaut Valley | 1020 | 2006 | 2014 |
| MC | McLeod Flat | 1280 | 2006 | 2014 |
| HS | Hetch Hetchy sign | 1400 | 2005 | 2014 |
| JM | Jackass Meadow | 2200 | 2008 | 2014 |
| ML | May Lake | 2774 | 2006 | 2014 |
| ME | Mammoth Edge | 3049 | 2006 | 2014 |
| HE | Hilgard Edge | 3095 | 2006 | 2014 |

**Table S2.** Results for the survival, conditional flower count, and lifetime fitness generalized linear models.

| Model | Fixed effect | Chisq | Df | p |
| --- | --- | --- | --- | --- |
| **Survival** |  |  |  |  |
|  | Generation | 2.3767 | 1 | 0.12 |
|  | Elevation | 9 | 1 | **0.0023** |
|  | Garden | 19 | 2 | **< 0.001** |
|  | Generation:Elevation | 3.3837 | 1 | **0.05** |
|  | Elevation:Garden | 17.0106 | 2 | **0.0002** |
| **Conditional flower count** |  |  |  |  |
|  | Generation | 0.2824 | 1 | 0.6 |
|  | Elevation | 0.6579 | 1 | 0.42 |
|  | Garden | 4.3163 | 2 | 0.12 |
|  | Generation:Elevation | 3.4067 | 1 | 0.06 |
| **Lifetime fitness** |  |  |  |  |
|  | Generation | 0.2868 | 1 | 0.59 |
|  | Elevation | 13.7092 | 1 | **0.0002** |
|  | Garden | 22.1648 | 2 | **< 0.001** |
|  | Generation:Elevation | 5.9871 | 1 | **0.014** |
|  | Elevation:Garden | 19.9721 | 2 | **< 0.001** |

**Table S3.** Results for the climate adaptation generalized linear models.

| Model | Fixed effect | Chisq | Df | p |
| --- | --- | --- | --- | --- |
| **Survival** |  |  |  |  |
|  | Generation | 37.4699 | 2 | < **0.001** |
|  | Group | 13 | 2 | **0.001** |
|  | Garden | 19 | 2 | < **0.001** |
|  | Generation:Group | 5.0131 | 2 | **0.08** |
|  | Generation:Garden | 1.6325 | 2 | 0.44 |
|  | Group:Garden | 16.867 | 4 | **0.002** |
|  | Generation:Group:Garden | 1.604 | 4 | 0.81 |
| **Lifetime fitness** |  |  |  |  |
|  | Generation | 3.1339 | 1 | 0.08 |
|  | Group | 9.6871 | 2 | **0.009** |
|  | Garden | 22.0662 | 2 | < **0.001** |
|  | Generation:Group | 6.8085 | 2 | **0.03** |
|  | Generation:Garden | 1.0636 | 2 | 0.59 |
|  | Group:Garden | 21.0913 | 4 | < **0.001** |
|  | Generation:Group:Garden | 0.7498 | 4 | 0.95 |

**Table S4.** Results for the ordinal phenology models.

| Model | Fixed effect | Chisq | Df | p |
| --- | --- | --- | --- | --- |
| **Low garden** |  |  |  |  |
|  | Generation | 0.0098 | 1 | 0.92 |
|  | Group | 9.436 | 2 | **0.008** |
| **Central garden** |  |  |  |  |
|  | Generation | 0.5017 | 1 | 0.49 |
|  | Group | 19.6913 | 2 | **< 0.001** |
| **High garden** |  |  |  |  |
|  | Generation | 4.3462 | 1 | **0.04** |
|  | Group | 0.5594 | 2 | 0.76 |

